# Tiling of large-scaled environments by grid cells requires experience

**DOI:** 10.1101/2025.02.16.638536

**Authors:** Blanca E. Gutiérrez-Guzmán, J Jesús Hernández-Pérez, Holger Dannenberg

**Affiliations:** Department of Bioengineering, George Mason University, 4400 University Dr., Fairfax, VA 22030

**Keywords:** Medial entorhinal cortex, grid cells, spatial navigation, spatial maps

## Abstract

Grid cells in the medial entorhinal cortex are widely believed to provide a universal spatial metric supporting vector-based navigation irrespective of the spatial scale of an environment. However, using single unit recordings in freely behaving mice, we demonstrate that spatial periodicity in grid cell firing is substantially disrupted when transitioning from a small to a large-scale arena when the scale ratio is larger than the scale ratio of successive grid modules. Remarkably, grid patterns reemerge with experience in the large-scale arena, suggesting that grid cells can learn to represent large-scale spaces with experience.

**Summary:** Scaling of grid maps is limited by the scale ratio of successive grid modules.

Grid maps cannot be sustained in novel large-scaled environments.

The recovery of the grid map requires multi-day experience.

## Main

Grid cells in the medial entorhinal cortex (MEC) contribute to a cognitive map of space by firing at multiple locations that fall on a hexagonal lattice. The spatially periodic spiking activity of grid cells^1,2^ is widely interpreted as a universal^3^, preconfigured^4,5^ and global metric for space^3^ in support of path integration^6–9^ and vector-based navigation^10^. However, more recent experimental data suggest that grid cells provide a local as opposed to a global metric by demonstrating that grid maps expand in novel environments^11^, appear sheared in asymmetrical environments^12,13^, overrepresent goal locations^14,15^, and linearly scale when an enclosure is compressed or stretched along one or two dimensions^16^. Furthermore, grid cells are organized anatomically and functionally into discretized grid cell modules along the dorso-ventral axis of the MEC, with a scale ratio of 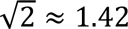 between modules^17^. While distortions and fragmentations of grid maps are permissible within the framework of a malleable universal map^18,19^ and are consistent with continuous attractor models^20–22^, it is unknown whether the hexagonal firing pattern of grid cells is universal across different environmental spatial scales. To answer this question, we recorded grid cells from mice freely foraging in three arenas, with scale ratios of the arena floors of 1, 1.33, and 1.66 relative to the first arena. Concretely, the floor dimensions of the small-, medium-, and large-size arenas were 45×45 cm^2^, 60×60 cm^2^, and 75×75 cm^2^. The scale ratios of 1.33 and 1.66 were selected because they are slightly smaller and larger than the scale ratio of 1.42 observed between successive grid cell modules, respectively. If spatial periodicity in grid cell firing were universal across spatial scales, we would expect either scaling^16^ or remapping ^2^ of grid maps across spatial scales, dependent on the stoichiometry of internal self-motion cues and external sensory cues in generating spatially periodic firing. If not, we would expect either permanent or transient disruptions of grid maps in the large-scale arena. On a typical recording day, a baseline session in the small arena was followed by a probe session in one of the scaled arenas (medium or large), followed by another baseline session in the small arena. To take into account the potential contribution of the walls to the spatial representation of the arena floor, we introduced high- and low-wall versions of the three enclosures.

To our surprise, spatial periodicity in grid cell firing was largely disrupted when an animal transitioned from a small-size arena with high walls to a large-size arena with low walls (scale ratio of arena floor: 1.66), quantified as a substantial reduction in the grid score (Fig. 1a,c,g; Extended Data Fig. 1a; Extended Data Table 1). Because walls can contribute to spatial periodicity in grid cell firing ^23^, we tested whether high walls in the large-scaled arena would rescue the disruption in spatial periodicity. We found that grid scores were also substantially reduced in the presence of high walls (Fig. 1b,d,h; Extended Data Fig. 1b; Extended Data Table 1), and it was not different to the low-wall arena (Fig. 1j; Extended Data Table 1). Next, we asked whether starting a recording day with the large-size arena allowed the formation of a grid map representation. However, the disruption of the grid map was invariant to the order of sequential arena exposures (Extended Data Fig. 3). Visual cues and encounters with walls of an environment can correct path intergration errors in the grid code^23–25^. We therefore quantified the contribution of wall height to grid scores. While we found a slight but statistically significant reduction in grid scores when mice transitioned from a small arena with high walls to a small arena with low walls (Fig. 1e,f,i-j and Extended Data Fig. 2), the effect was small relative to the almost complete disruption observed in the enlarged arenas (Fig. 1e,f,i-j; Extended Data Table 1). Low walls thus do not explain the substantial reduction in spatial periodicity associated with a large-scaled arena floor. Taken together, these results demonstrate that grid maps depend on the spatial scale of the arena. This finding seems at odds with previous reports that spatial periodicity in grid cell firing is largely invariant across environments and with the finding that grid maps scale linearly with changes in the arena size ^16^. To reconcile our findings with previous reports on grid map scaling, we tested the hypothesis that grid maps can represent a scaled environment if the scale ratio is smaller than the scale ratio of successive grid modules.

**Fig. 1:**
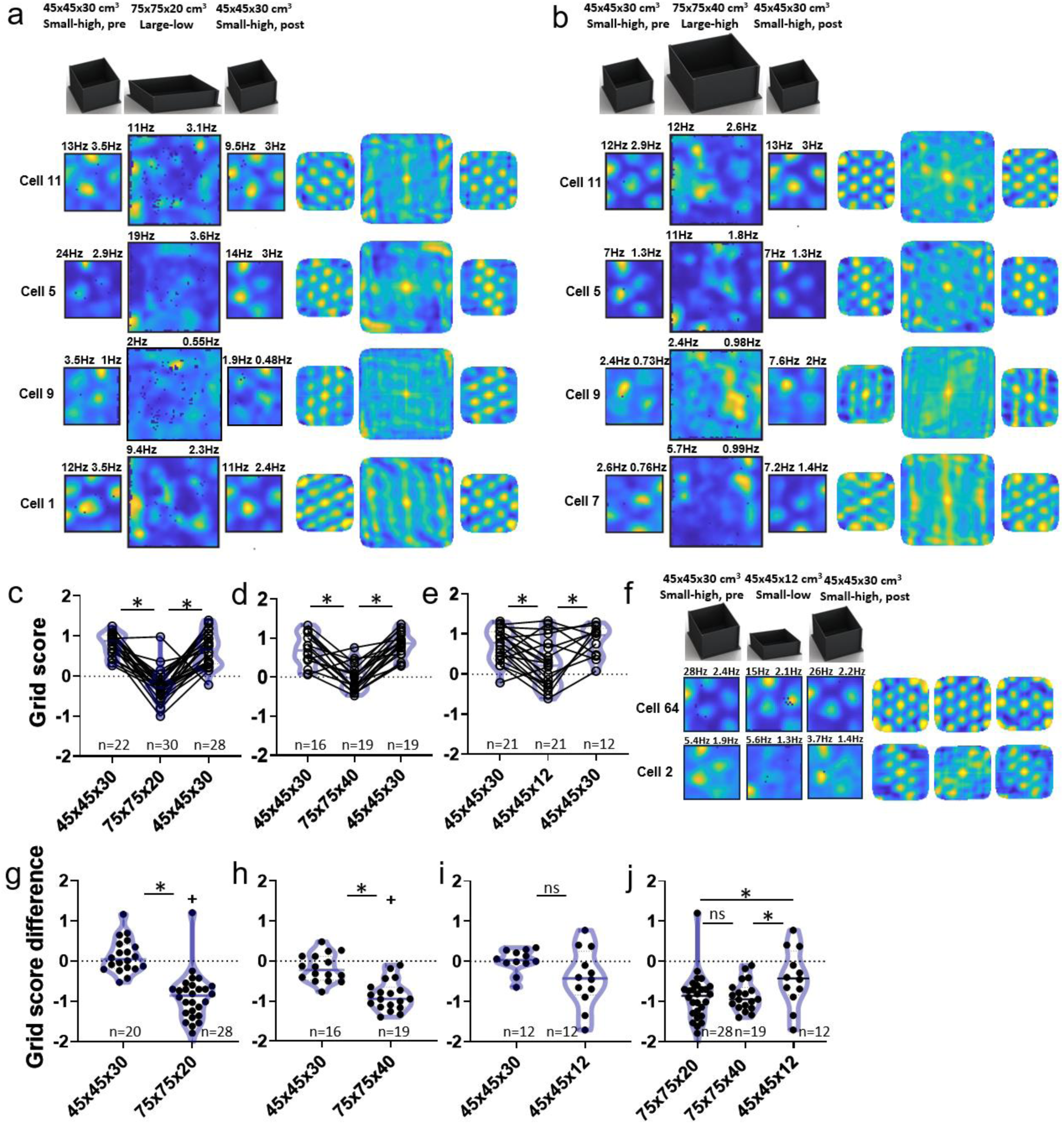
Spatial periodicity in grid cell firing is disrupted in large-scaled environments. **(a)** Left, firing rate maps of four grid cells (cell 11, cell 5, cell 9, and cell 1 from mouse #1) recorded sequentially in a small-size arena with high walls (45×45×30 cm^3^, Small-high, pre), a large-size arena with low walls (75×75×20 cm^3^, Large-low), and again in the previous small-size arena (Small-high, post). Numbers above the map show peak and mean rates. Right, spatial autocorrelograms of the rate maps. Yellow and blue colors indicate high and low values, respectively. Data on cells 11 and 5 shown in **a** and **b** were recorded on different days. (**b**) Data on four grid cells (cell 11, cell 5, cell 9, and cell 7 from mouse #1) recorded in the sequence “Small-high, pre; Large-high; Small-high, post”, where “Large-high” indicates a large environment with high walls (75×75×40 cm^3^). Data presented in the same way as in **a**. (**c-e)** Comparison of grid scores across different arenas: 45×45×30; 75×75×20; 45×45×30. Violin plots show the distribution of grid scores for each arena. **\***, p < 0.05, Tukey’s multiple comparisons test. **(f)** Data on two grid cells recorded in the sequence “Small-high, pre; Small-low; Small-high, pre” (cell 64 and 2 from mouse #1 and #2). Data presented in the same way as in **a** and **b**. **(g-j**) Data points show the difference in grid scores relative to the second small arena (45×45×30 cm, post) within a sequence. (**g-i**, Wilcoxon matched-pairs Signed Rank Test: 45×45×30 vs. 75×75×20, W = -208, p <0.0001; 45×45×30 vs. 75×75×40, W = -124, p = 0.0004; 45×45×30 vs. 45×45×12, W= -23, p = 0.3804, respectively; **j**, Kruskal-Wallis statistic = 11, p = 0.0041; Dunn’s multiple comparisons test). **\***, p (difference between groups) < 0.05; +, p (difference from zero) < 0.05. ns = not significant. n = number of cells. See Extended Data Table 1 for statistical analysis and the number of cells in each comparison.

Indeed, the grid cell representation was maintained in the medium-scaled arena regardless of the wall (Fig. 2a-e, Extended Data Fig. 4 and Extended Data Table 2). This result demonstrates that the capability to maintain a grid representation in scaled arenas depends on the scale ratio (Fig. 2i-l, and Extended Data Table 2). Notably, an intermediate scaling step to the medium-scaled arena before transitioning to the large-scaled arena could not rescue the disruption of the hexagonal firing pattern in the large-scaled arena (n = 4 cells) (Extended Data Fig. 5). In concordance with a previous report ^16^, grid spacing scaled with the increase in arena floor size irrespective of the wall height (Fig. 2f-h). However, grid cells showed significant remapping when transitioning to the medium-scaled maze with low walls (Fig. 2m1,n) but not when transitioning to the medium-scaled maze with high walls (Fig. 2m2,o). These data are in alignment with previous reports that grid maps are anchored to external sensory cues such as boundaries^2,12,13,26^.

**Fig. 2:**
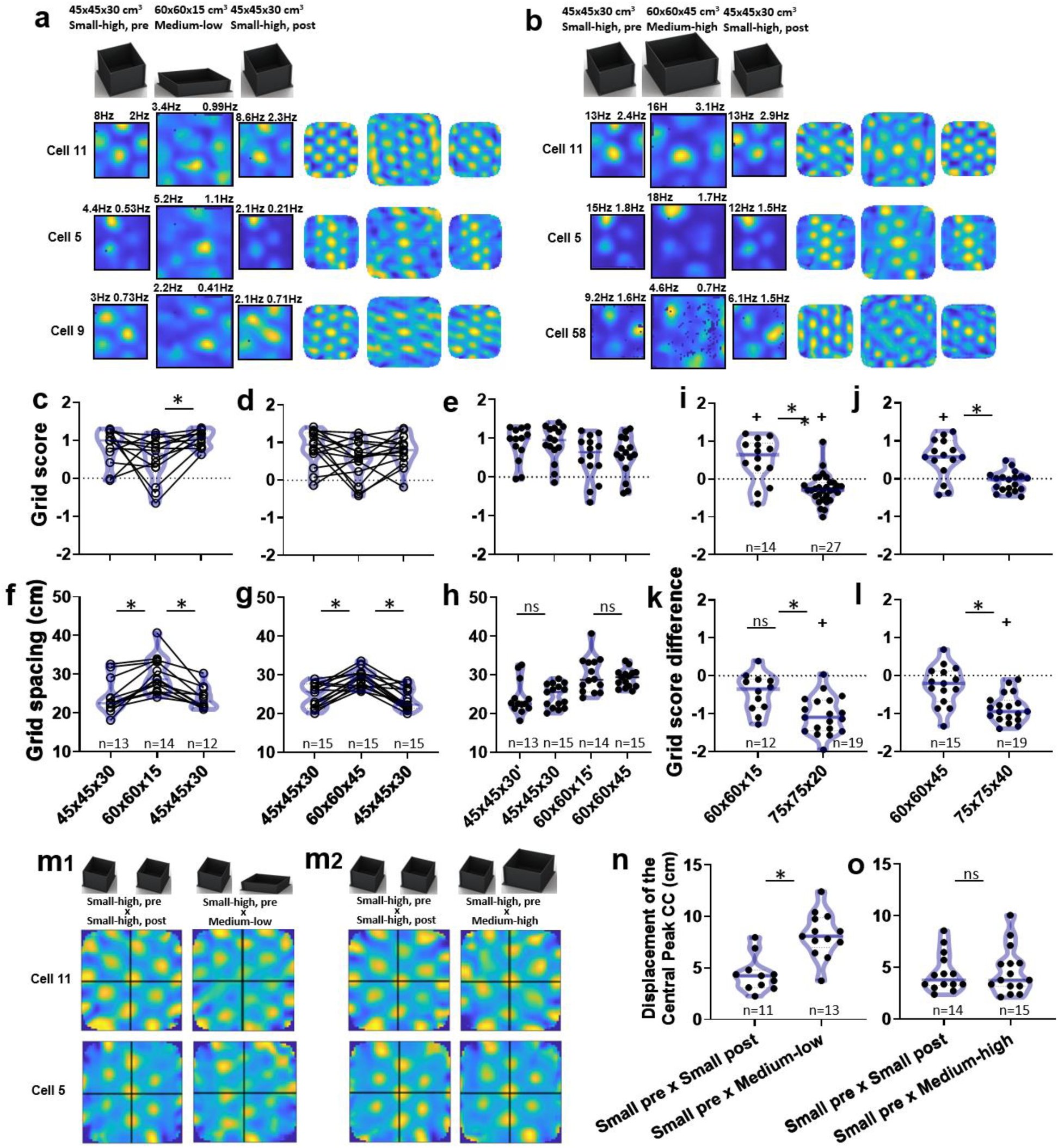
Spatial periodicity in grid cell firing is maintained in medium-scaled environments. **(a)** Left, firing rate maps of three grid cells (cell 11, cell 5, cell 9 of mice#1) recorded sequentially in a small-size arena with high walls (45×45×30 cm^3^, Small-high, pre), a medium-size arena with low walls (60×60×15 cm^3^, Medium-low), and again in the previous small-size arena (Small-high, post). Numbers above the map show peak and mean rates. Right, spatial autocorrelograms of the rate maps. Yellow and blue colors indicate high and low values, respectively. (**b**) Left, firing rate maps of three grid cells (cell 11, 5 and 58 from mice#1 and #2) recorded in the sequence “Small-high, pre; Medium-high; Small-high, post”, where “Medium-high” indicates a medium-size environment with high walls (60×60×45 cm^3^). Data presented in the same way as in **a**. (**c-e)** Comparison of grid scores across different arenas. Violin plots show the distribution of grid scores for each arena (c) *, p < 0.05, Tukey’s multiple comparisons test. **e**, Grid scores recorded in medium arenas with low (60×60×15’) and high walls (60×60×45) after transitioning from small-size arenas (45×45×30’ and 45×45×30, respectively; same cells as in (**c-d**) (Kruskal-Wallis statistic = 7.261, p= 0.064). (**i-j**) Comparison of grid scores between the population of grid cells recorded using the medium-size and large-size mazes with low (**i**) and high (**j**) walls (Mann Whitney test: U = 43, p=0.0001; U = 47, p=0.0006, respectively). **(f-h)** Comparison of grid spacing across different arenas. Violin plots show the distribution of grid spacing for each arena (f-g), **\***, p < 0.05, Tukey’s multiple comparisons test. h, (Kruskal-Wallis statistic = 19.52, p= 0.0002). Data shown as in **c-e**. **(k-l**) Data points show the difference in grid scores relative to the second small arena (45×45×30, post) within a sequence (Mann Whitney test: U = 30, p=0.002; U = 51, p=0.001, respectively). (**m**) Data on two grid cells showing the spatial crosscorrelograms (cc) computed from grid maps associated with different arenas. Crosscorrelograms computed from grid maps associated with pre and post sessions recorded in the same arena (Small-high, pre; Small-high, post) serve as controls. (**n,o**) Population data on the displacements of the peaks in the spatial cc computed from the rate maps of the “small-high, pre” and “Medium-low” arenas (**n**) and the “small-high, pre” and “Medium-high” arenas (**o**) (Mann Whitney test: U = 13.50 p=0.0003; U = 104, p=0.9742, respectively). Data on the displacements of peaks in crosscorrelograms computed from “pre” and “post” sessions in the small arenas are shown for comparison. **\***, p (difference between groups) < 0.05; +, p (difference from zero) < 0.05. ns = not significant. n = number of cells. See Extended Data Table 2 for statistical analysis and the number of cells in each comparison.

Next, we asked whether the disruption of spatially periodic firing of grid cells observed with transitions to large-scaled arenas was permanent or could revert with experience. To answer this question, we continued recording grid cell activity in the same mice transitioning from the small-size to the large-scaled arena up to ∼2 months (14 sessions in two mice). Intriguingly, a spatially periodic firing emerged in the large-scaled arena after several days of training (Fig. 3a-e,h-i,k) irrespective of the wall height in the large-scaled arena (Fig. 3e, Extended Data Fig. 6 and Extended Data Table 3). On average, spatially periodic firing was observed after 7.4 recording sessions (Fig. 3k2, Extended Data Fig. 7 and Extended Data Table 3).

**Fig. 3:**
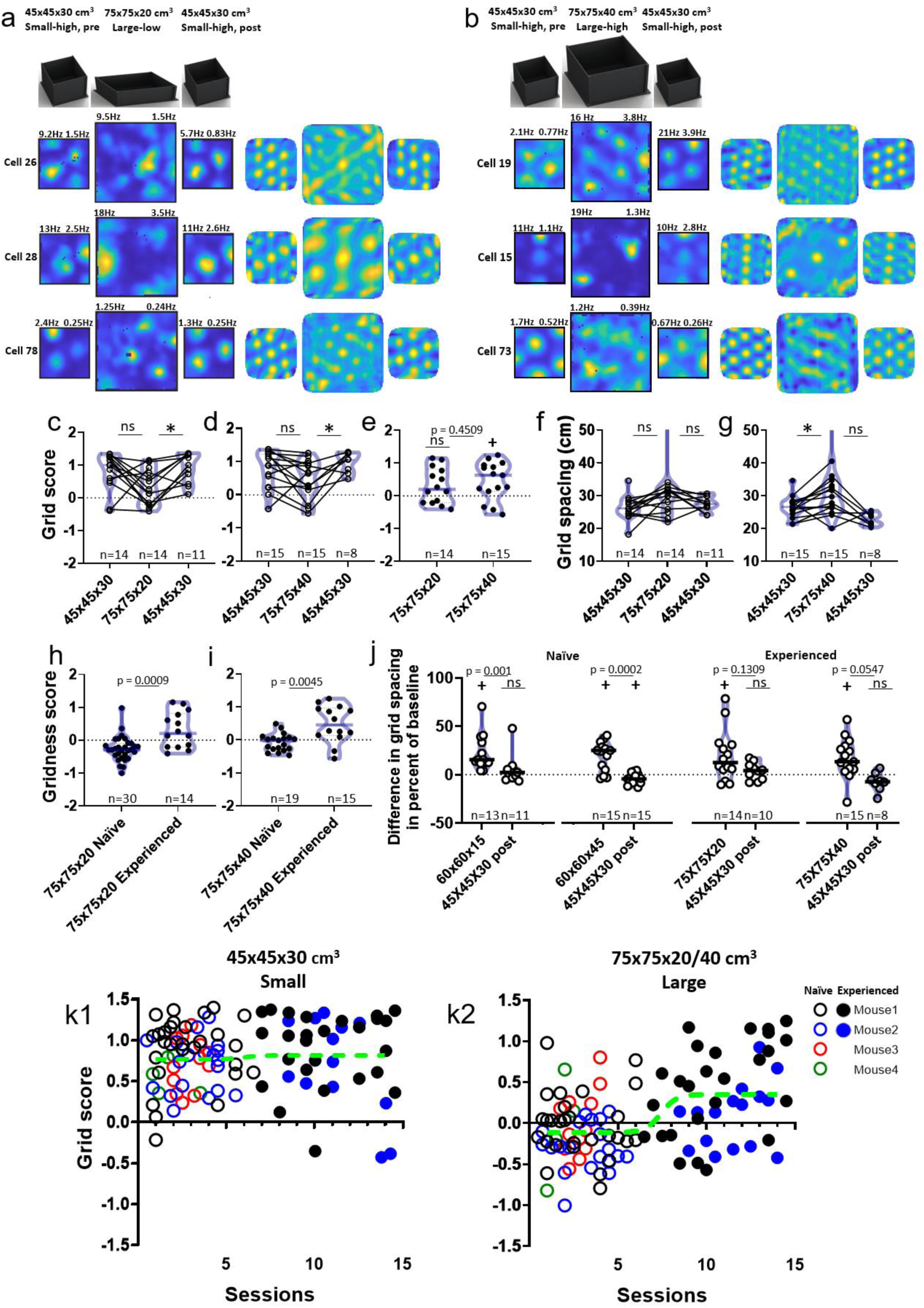
Grid maps in large-scaled environments are experience-dependent. **(a)** Left, firing rate maps of three grid cells after gaining experience (cell 26, cell 28 and cell 78 of mice #1 and #2) associated with the exploration of a small environment with high walls (45×45×30 cm^3^; small-high, pre and post) and a large-scaled environment with low walls (75×75×20 cm^3^; Large-low). Numbers above the rate maps show peak and mean firing rates. Right, spatial autocorrelograms of the rate maps. Yellow and blue colors indicate high and low values, respectively. **(b)** Left, firing rate maps of four grid cells after gaining experience (cell 19, cell 15, cell 22 and cell 73 of mice #1 and #2) associated with the exploration of a small environment (small-high, pre and post) and a large-scaled environment with high walls (75×75×40 cm^3^; Large-high). **(c-e)** Grid scores across different arenas after gaining experience. Violin plots show the distribution of grid scores for each arena (c-d) *, p < 0.05, Tukey’s multiple comparisons test. e, Mann Whitney test: U = 87, p = 0.4509; Wilcoxon Signed Rank Test; 75×75×20 Experienced, W = 51, p = 0.1189, 75×75×40 Experienced, W = 84, p = 0.0151). Note that grid scores associated with the large-scaled arena show little to no reduction relative to the grid scores associated with the small-size arena, irrespective of the presence or absence of walls. **(f-g)** Comparison of grid spacing across different arenas (same cells as in **c-e**) presented in the same way as in **c-e**. **(g)** *, p < 0.05, Tukey’s multiple comparisons test. **(h,i)** Data on grid scores before and after gaining experience by multiple repetitions over multiple days in exploring the large-scaled arena (data on low and high walls combined) (Mann Whitney test: U = 82, p = 0.0009; U = 62, p = 0.0045 respectively). **(j)**. Percentage change in grid spacing relative to the spacing measured when animals explored the small-size arena with high walls (small-high, pre) at the beginning of each sequence (Wilcoxon matched-pairs signed rank test: 60×60×15 Naïve vs. 45×45×30 Post, W = -66, P = 0.001; 60×60×45 Naïve vs. 45×45×30 Post, W= -116, p = 0.0002; 75×75×20 Experienced vs. 45×45×30 Post, W =-31, p = 0.1309; 75×75×40 Experienced vs. 45×45×30 Post, W = -28, p = 0.0547). **(k)** Grid scores before (open circle) and after (filled circle) gaining experience plotted as a function of the number of sessions mice explored the large-scaled arena. Different colors indicate data from four different mice. **k**1, data shows how grid scores measured in mice exploring the small-size arena change over multiple recording sessions performed at different days; **k**2, data show how grid scores measured in mice exploring the large-scaled arena change over multiple recording sessions performed at different days. Using a sigmoid fit function, the half-maximal increase was observed after 7.4 (CI: 5 – 13.5) recording sessions. *, p < 0.05 (difference between groups); +, p < 0.05 (difference from zero); ns = not significant. n = number of cells. See Extended Data Table 2 for statistical analysis and the number of cells in each comparison.

After mice gained experience (Experienced) in the large-scaled arena, grid spacing increased when transitioning from a small-scale arena regardless of the height of the walls (Fig. 3f-g,j and Extended Data Table 3). However, this increase in spacing was not larger than the increase after transitioning to a medium-size arena suggesting that the extent to which grid maps can scale is limited (Fig. 3j). Taken together, our data demonstrate that the spatial periodicity in grid cell firing in large-scaled environments depends on the animal’s level of familiarity with the environment.

To summarize, we here show for the first time that grid cells’ spatially periodic firing is substantially disrupted in a large-scaled arena, challenging the supposition that the spatial code by individual grid cells is universal across all environments. Remarkably, the disruption in the hexagonal firing pattern was only observed if the scale ratio was larger than the scale ratio of 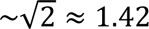 observed in the geometric progression of spacing across grid cell modules, suggesting that scaling of grid maps is constrained by the spatial scale of the module. While the underlying factors determining the exact value of the scale ratio of 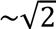 between modules are unknown, the modular architecture may enable a computationally efficient multiscale representation of space or spatial trajectories^27,28^. According to these scale-space theories, each module represents a specific spatial scale of the environment, thereby constraining the extent to which grid spacing can change in response to environmental manipulations. Within these constraints, and consistent with previous studies^2,16^, grid maps can either scale, remap, or both, with high walls favoring scaling and low walls favoring remapping (Fig. 2m-o).

Intriguingly, the spatially periodic firing of grid cells reappeared after animals gained experience over multiple sessions and days in exploring the large-scaled environment (Fig. 3a-e,h-i). Previous studies have shown that grid maps can be reshaped by experience^11,16,29,30^. However, in these previous studies, grid maps were present at all times across environments. Here, in contrast, we show that grid maps are substantially disrupted in large-scaled arenas and reappear with experience. Interestingly, a recent developmental study investigating grid cell firing showed that spatially periodic firing in rats raised in an opaque spherical environment appeared only after multiple days of experience in an arena with clear geometric boundaries^31^. Applying a developmental timescale of weeks to months, the authors interpreted the delay of multiple days as evidence for a preconfigured grid map. However, the timeline is similar to how long it takes for grid cells in adult mice to learn the representation of a large-scaled space, suggesting that the same mechanism underlie both phenomena. It would be a worthwhile endeavor for future studies to investigate the mechanisms underlying the disruption and recovery of grid cell representations in large-scaled environments. One possibility is that the transient disruption of spatially periodic firing in large-scaled arenas results from a mismatch between the scaled external sensory cues and internal self-motion cues. Alternatively, mice may engage smaller grid cell modules to represent space at a finer spatial resolution when the space becomes more familiar.

## Methods

### Animals

All experimental procedures were performed in accordance with the regulations stated in the Guide for the Care and Use of Laboratory Animals by the National Research Council and approved by the Institutional Animal Care and Use Committee of George Mason University (Protocol No.: 0501, approved on January 24, 2022).

Four male mice (*Mus musculus*), 4-6 months old (two wild type, C57BL/6, and two heterozygous transgenic ChAT-Cre^tg/wt^, B6;129S6-Chat^tm1(cre)Lowl^/MwarJ; The Jackson Laboratory) were used. Prior to surgery, mice were housed in Plexiglas cages together with their siblings. After surgery, mice were housed individually in large cages on a12-h reversed light/dark cycle with food and water *ad libitum*. Cages contained a spherical treadmill for physical exercise and as environmental enrichment. All mice were handled and habituated to the experimenter and testing room prior to the start of behavioral experiments and electrophysiological data acquisition. To increase the motivation in mice to explore the arena, bits of cereal were randomly placed on the arena floor, and mice were mildly food-deprived starting one day before the start of behavioral experiments so that their body weight was >=95% the weight of their littermates with *ad libitum* access to food.

### Surgery and electrodes

Microdrives were implanted as previously reported^24,32^. Briefly, the mice were anesthetized with isoflurane (4% induction, 1-2% maintenance) and placed in a stereotaxic apparatus. Physiological body temperature (37 °C) was monitored and maintained during the surgery via a heating pad and a homeothermic monitoring system. A craniotomy was performed, and mice were chronically implanted with a microdrive (Axona, St. Albans, UK) containing 4 movable tetrodes of twisted 17-µm coated platinum wires (impedance adjusted to 180-250 kOhm), targeting the superficial layers of the left MEC at the following stereotaxic coordinates: 3.4 mm lateral to the midline and 0.5 mm anterior to the transverse sinus, angled at a polar angle of six degrees. The tetrode tips were lowered 1 mm from the brain surface. A stainless-steel screw implanted in the skull above the cerebellum contacting the dura mater served as a ground electrode. The microdrive was secured to the skull using four additional anchoring screws and dental cement. For an additional experimental purpose not reported in this study, two mice were injected with an adeno-associated virus coding for the calcium indicator GCaMP8s into the medial septum and implanted with an optical fiber placed above the medial septum. No differences were found in those two mice with respect to the electrophysiological and behavioral metrics used in this study. Mice were allowed to recover from surgery for one week before the start of behavioral experiments and data acquisition.

### Behavioral task

Extracellular recordings of action potentials (spikes) in the MEC were carried out while mice randomly foraged for scattered pieces of cereal bits (Froot Loops, Kellogg Company, Battle Creek, MI, USA) in different sizes of black open field environments. Recording sessions lasted until animals had sampled a sufficiently large area of the open field, typically 20 – 45 mins. One visual cue card (a white triangle) was placed on one of the walls of the open field enclosure. All enclosures were centered in the same location on a black floor mat in the same room with the same distal cues. To test the effect of scaling the arena size on grid cells, data were acquired while mice explored a sequence of square arenas with different sizes, namely a small maze, 45 x 45 cm^2,^ with either 30 cm high or 12 cm low walls (Small-high or Small-low, respectively); a medium maze, 60 x 60 cm^2^, with 45 cm high or 15 cm low walls (Medium-high or Medium-low, respectively); and a large maze, 75 x 75 cm^2^, with 45 cm high or 20-cm low walls (Large-high or Large-low, respectively). Four main sets of sequences of different arenas were used through multiple sessions: 1) Small-high to Large-low to Small; 2) Small to Large-high to Small; 3) Small to Medium-low to Small; and 4) Small to Medium-high to Small. Each sequence started and ended with the small arena with high walls (Small Pre and Small Post, respectively). In between sessions, mice were allowed to rest in their home cage in the same room. The interval between individual sessions was 20-30 min. Between sessions, the floor of the open field was cleaned with 70% isopropanol. Some different sets of sequences were evaluated (Extended data Fig. 3,4c,5,6).

### Neural recordings

If not described differently, neural recordings and analysis of grid cells was performed as described previously^24^. Tetrodes were lowered daily in increments of 50 micrometers until multiunit spiking activity was detected, indicating that tetrodes had reached the superficial layer in the MEC. The presence of grid cells was assessed by shorter ∼20 min screening sessions using the small or medium-size maze with high walls. If grid cells were present, the full set of arena sequences was employed. As long as at least one grid cell was present in the small maze, recording sessions continued for several days. If no grid cells were present anymore, the electrodes were advanced ∼25 micrometers to find new grid cells and similar sessions were evaluated.

Neural signals were pre-amplified by unity-gain operational amplifiers located on each animal’s head (head stages) and AC-wire-coupled to an Axona recording system (Axona Limited, St. Albans, UK). Spikes were threshold-detected and recorded at 48 kHz. The position and head direction of mice were captured using a ceiling-mounted video camera to record the position of invisible infrared LEDs on the animal headstage (at 50 Hz) and tracking software (Axona, UK).

Spikes were clustered into single units using spike clustering software (DacqUSB/Tint, Axona, St. Albans, UK). Further offline analyses were performed using Matlab (MathWorks, Natick, MA, USA).

### Histology

After data collection, animals were anesthetized deeply with isoflurane and a transcranial perfusion with DPBS 1X + 10% buffered formalin was performed. To visualize tetrode track and MEC recording sites, sagittal brain sections (40-50 µm) were sliced using a vibratome (Leica VT1000S) and stained with a Cresyl violet solution.

### Analysis of single unit spiking activity

#### Spatial firing rate maps

The occupancy-normalized spatial firing rate maps were generated by dividing the open-field environment into 1.5 cm spatial bins. For each spatial bin, an occupancy-normalized firing rate was computed by dividing the total number of spikes falling into that bin by the total time the animal spent in that bin. Spatial rate maps were then smoothed by a 2 cm wide two-dimensional Gaussian kernel.

#### Grid score

The grid score was used as a measure of hexagonal symmetry of grid fields as described previously^33^. Briefly, we first computed the two-dimensional autocorrelogram of the spatial rate map, extracted a ring that encased the six peaks closest to the center peak excluding the central peak and computed correlations between this modified spatial autocorrelogram and rotated versions of itself after correction of elliptical eccentricity. The correlation was calculated for each 3-degree rotation of the donut to itself. The gridness score is the difference between the correlation at the minimum peak of 60 or 120 degrees and the maximum trough at 30, 90, or 150 degrees.

#### Grid spacing

The grid spacing was calculated from the spatial autocorrelogram by identifying the center of mass for each of the six closest fields and calculating the distance of the fields peak from the center peak. Because the autocorrelogram is symmetric, the average distance of the three consecutive peaks to the center was calculated.

#### Spatial crosscorrelation

The spatial crosscorrelation was calculated to evaluate the similarity of grid maps for each grid cell across different sessions or environments. To match the matrix size of the spatial firing rate map of small and medium-size maze, a reduction of matrix size of medium-maze was achieved by increasing the bin size proportional to the scaling factor of maze enlargement [bin size (1.5 cm^2^) * enlargement factor (1.333)]. To compute the spatial correlation, the spatial bins of the firing rate maps were transformed into a single vector. The spatial correlation was then computed as the Pearson’s correlation coefficient between two spatially binned firing rate vectors. From the cross-correlogram, a similar grid map between two sessions will show a triangular grid pattern and the central peak close to the origin of the crosscorrelogram, and while different grid map will show a displacement of the central peak from the origin, but a preserved or distorted grid pattern could appear. The distance to the peak nearest to the center of the cross-correlogram was used as an indicative of similarity of grid map.

### Statistical analysis

Results of statistical analyses are reported in the Extended Data Tables 1, 2, and 3. Briefly, Mixed model fixed effects (Type III) analysis for repeated measures and post-hoc Tukey’s multiple comparisons tests were performed. (“Repeated measures ANOVA cannot handle missing values. We analyzed the data instead by fitting a mixed model as implemented in GraphPad Prism 8.0. This mixed model uses a compound symmetry covariance matrix, and is fit using Restricted Maximum Likelihood (REML). In the absence of missing values, this method gives the same P values and multiple comparisons tests as repeated measures ANOVA. In the presence of missing values (missing completely at random), the results can be interpreted like repeated measures ANOVA”). In addition, Nonparametric statistical tests such as the Mann Whitney test and the Wilcoxon’s signed-rank test were applied using GraphPad Prism software (version 10.4.1).

## Code accessibility

https://github.com/dannenberglab/

## Acknowledgements

This work was supported by the National Institute of Neurological Disorders and Stroke and the National Institute of Aging of the National Institutes of Health, grant numbers R00NS116129 and R21AG087912. The authors have no conflicts of interest.

## Extended Data Figures

**Extended Data Fig. 1:**
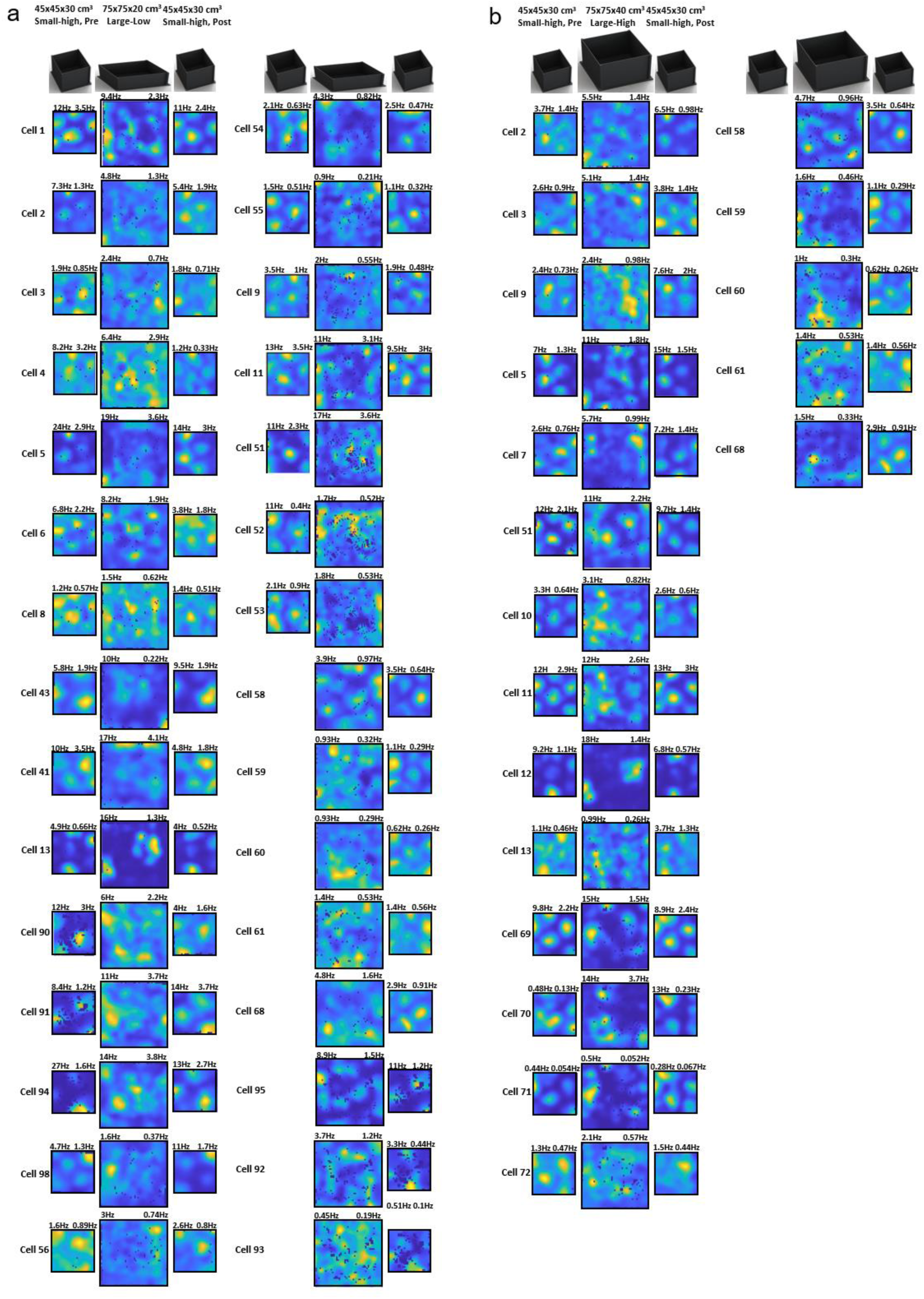
Firing rate maps of all grid cells recorded in naïve mice before gaining experience on the large-scaled maze. Mice explored a sequence of arenas within one recording day. Arenas included a small-size arena with high walls (45×45×30 cm^3^, Small-high), and a large-scaled version of the arena with either **a)** low walls (75×75×20 cm^3^, Large-low) or **b)** high walls (75×75×40 cm^3^, Large-high). These cells are part of the statistical analysis in Fig. 1. Cells 58, 59, 60, 61, and 68 from large-size arena with low walls and high walls were recorded the same day with the sequence (75×75×40 - 75×75×20 - 45×45×30). The numbers above the map show peak and mean firing rates. Yellow and blue colors indicate high and low values, respectively.

**Extended Data Fig. 2:**
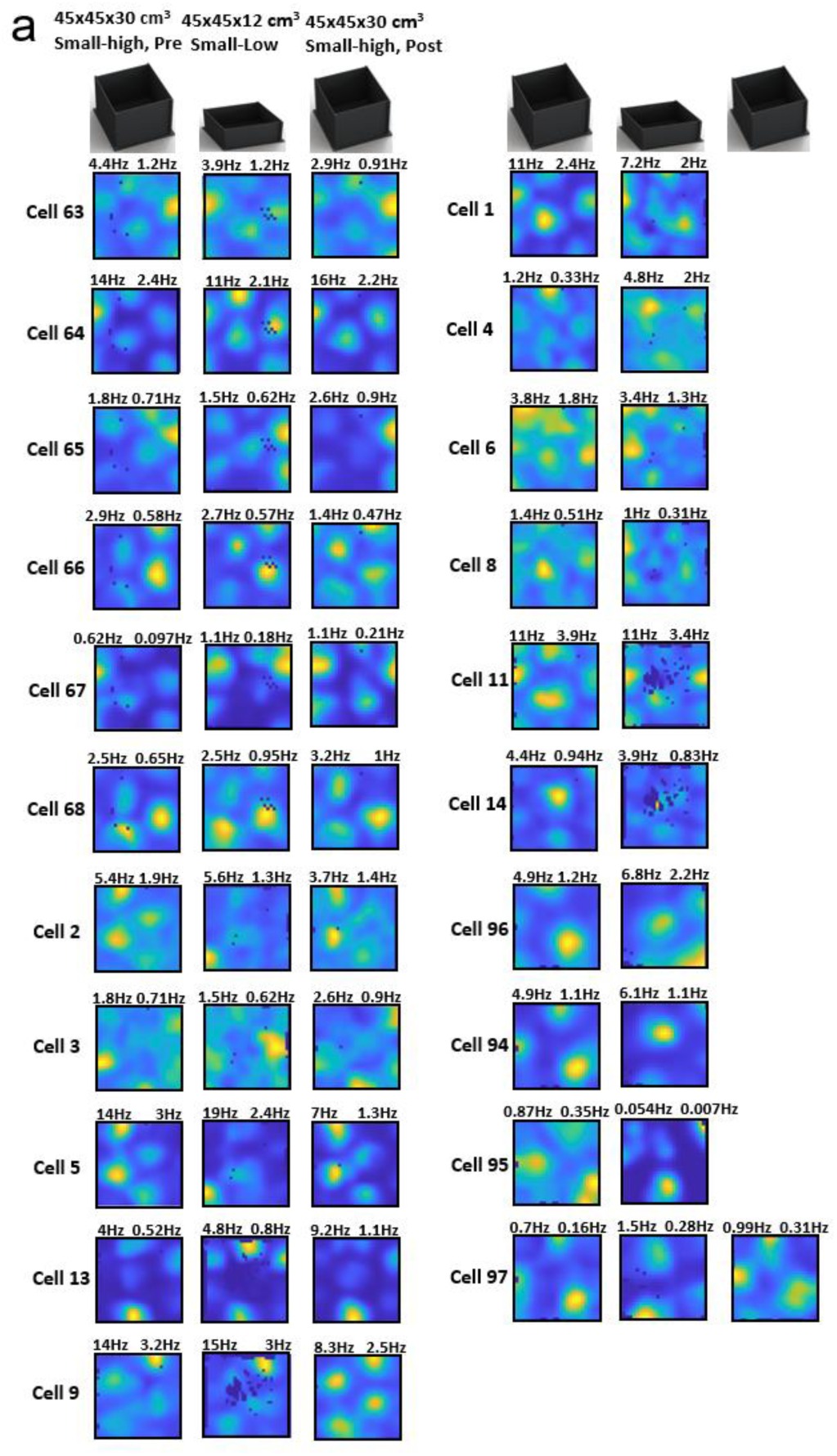
Firing rate maps of all grid cells, where mice explored sequentially small-size arenas with different wall heights (45×45×30 cm^3^, Small-high; and 45×45×12 cm^3^, Small-low). These cells are part of the statistical analysis performed on data shown in Fig. 1e,j. The numbers above the maps show peak and mean firing rates. Yellow and blue colors indicate high and low values, respectively.

**Extended Data Fig. 3:**
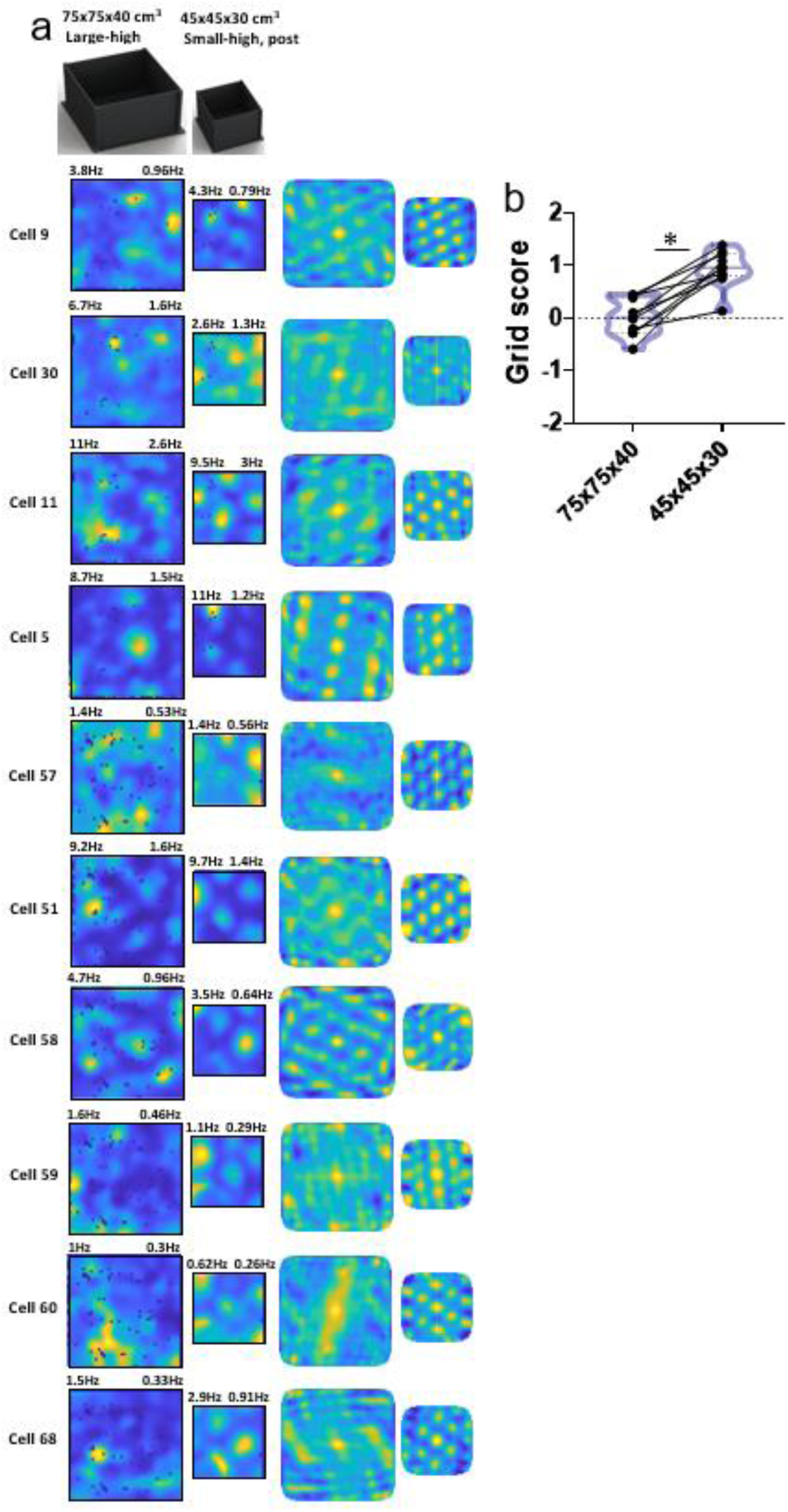
Grid maps are disrupted in large-size arenas even if mice had not previously explored a small-size arena on the day of the recording. **a)** Firing rate maps and spatial autocorrelograms of ten grid cells recorded sequentially in a large-size arena with high walls (75×75×40 cm^3^, Large-high) and a small-size arena (45×45×30 cm^3^, Small-high, post). Numbers above the map show peak and mean rates. Yellow and blue colors indicate high and low values, respectively. **b)** Comparison of grid scores across the two arena sizes. Violin plots show the distribution of grid scores associated with each arena. Pairwise comparison (Wilcoxon matched-pairs signed rank test: W = -55.00, *, p < 0.0020, n = 10).

**Extended Data Fig. 4:**
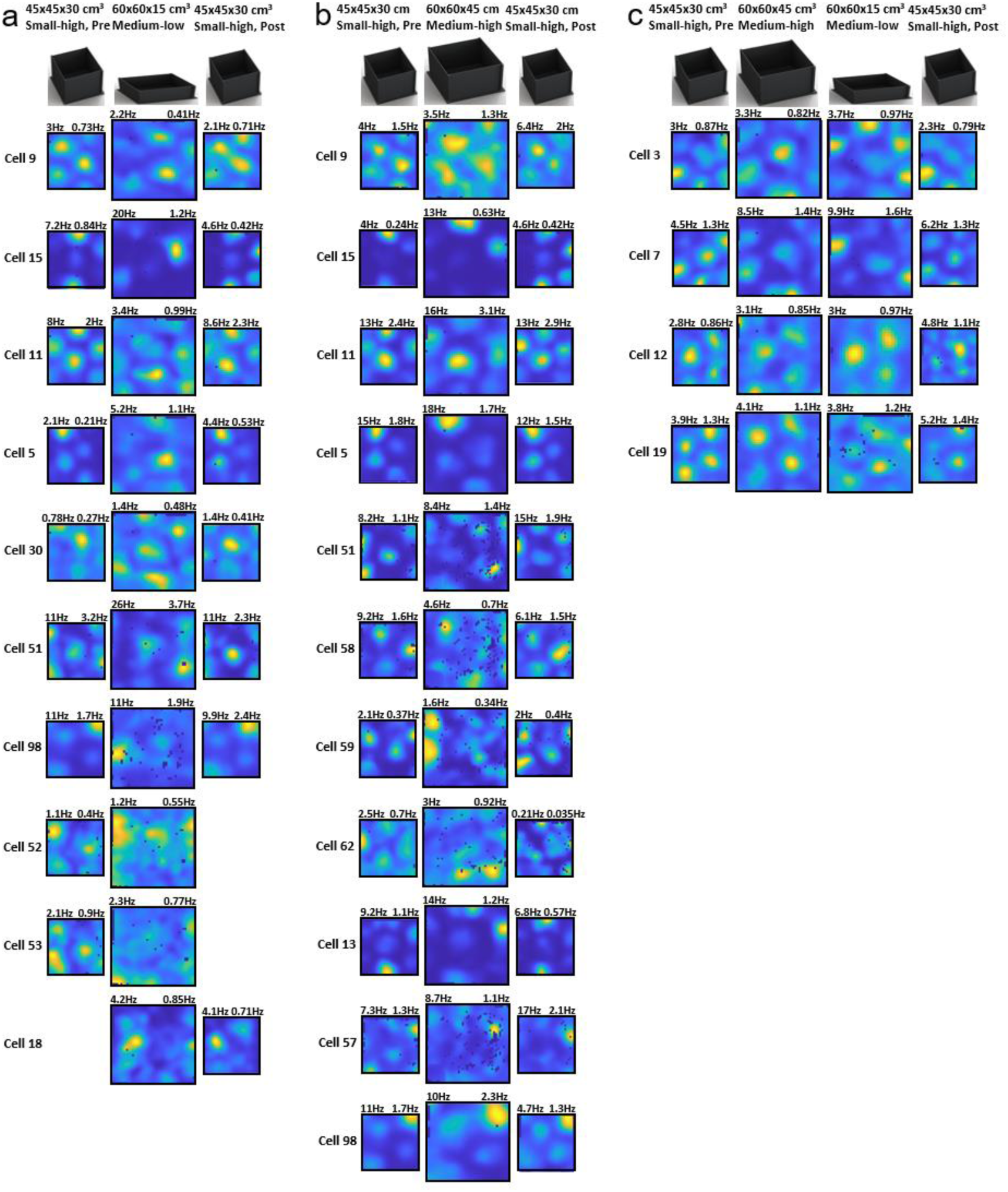
Firing rate maps of all grid cells recorded in naïve mice before gaining experience on the medium-scaled maze. Mice explored different sets of sequences of arenas within one recording day. Arenas included a small-size arena with high walls (45×45×30 cm^3^, Small-high) and a medium-size arena with either **a)** low walls (60×60×15 cm^3^, Medium-low) or **b)** high walls (60×60×45 cm^3^, Medium-high). **c)** Data on cells 3, 7, 12 and 19 belong to a sequence with four arenas that were included in **a** and **b**. Numbers above the map show peak and mean rates. Yellow and blue colors indicate high and low values, respectively.

**Extended Data Fig. 5:**
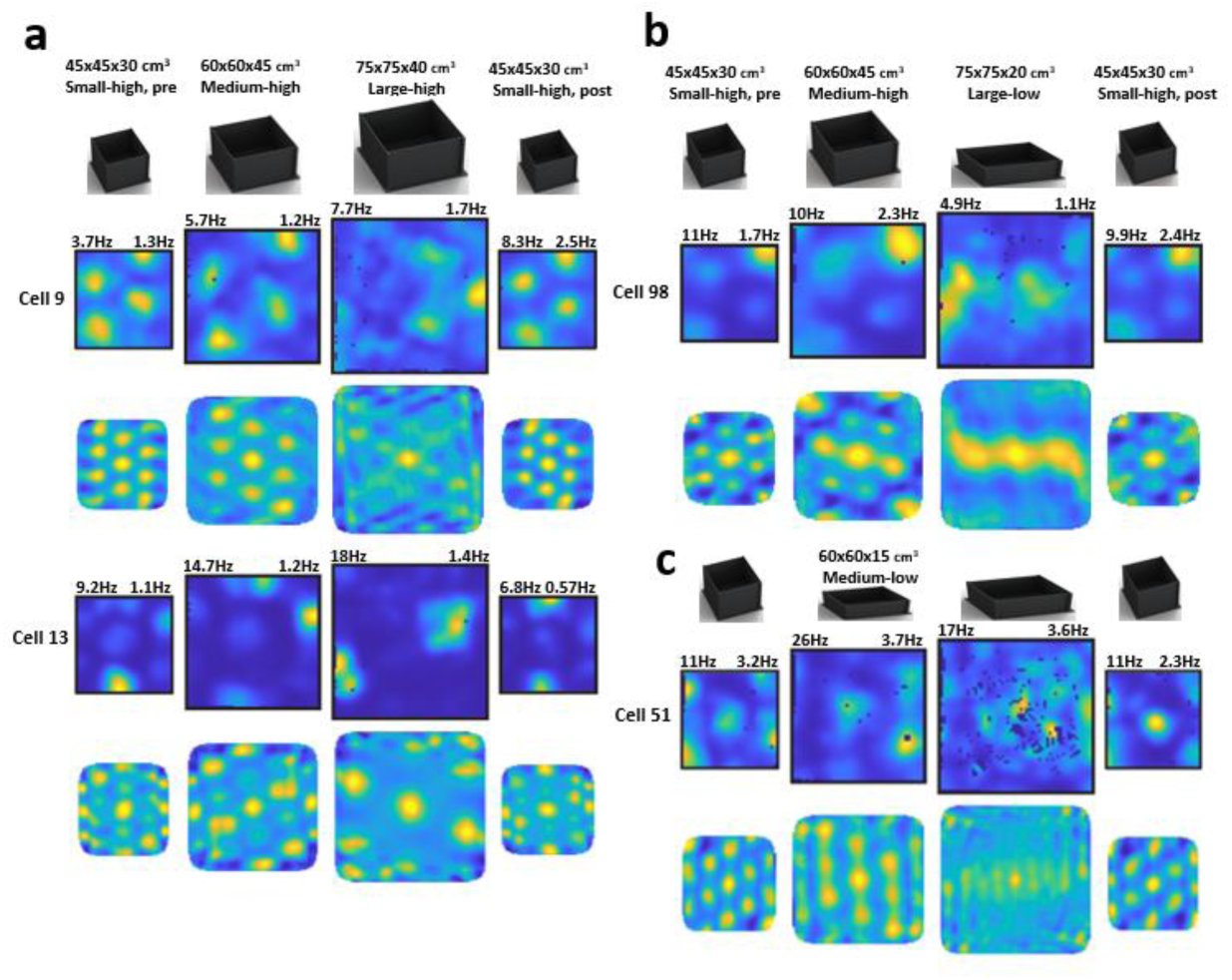
An intermediate scaling step before transitioning to the large-scaled arena fails to rescue the disruption of grid maps. **a)** Firing rate maps and spatial autocorrelograms of two grid cells (cell 9 and cell 13) recorded subsequently in a small-scale arena with high walls (45×45×30 cm^3^, Small-high, pre), a medium-scaled arena will high walls (60×60×45 cm^3^, Medium-high), a large-scaled arena with high walls (75×75×40 cm^3^, Large-high), and the small-scale arena (Small-high, post). Numbers above the map show peak and mean firing rates. Yellow and blue colors indicate high and low values, respectively. **b)** Firing rate maps of one grid cell (cell 98) recorded in the sequence “Small-high, pre; Medium-high; Large-low and Small-high, post”, where “Large-low” indicates a large environment with low walls (75×75×20 cm). Data presented in the same way as in **a**. **c)** Firing rate maps of one grid cell (cell 51) recorded in the sequence “Small-high, pre; Medium-low; Large-low and Small-high, post”. Data presented in the same way as in **a** and **b**.

**Extended Data Fig. 6:**
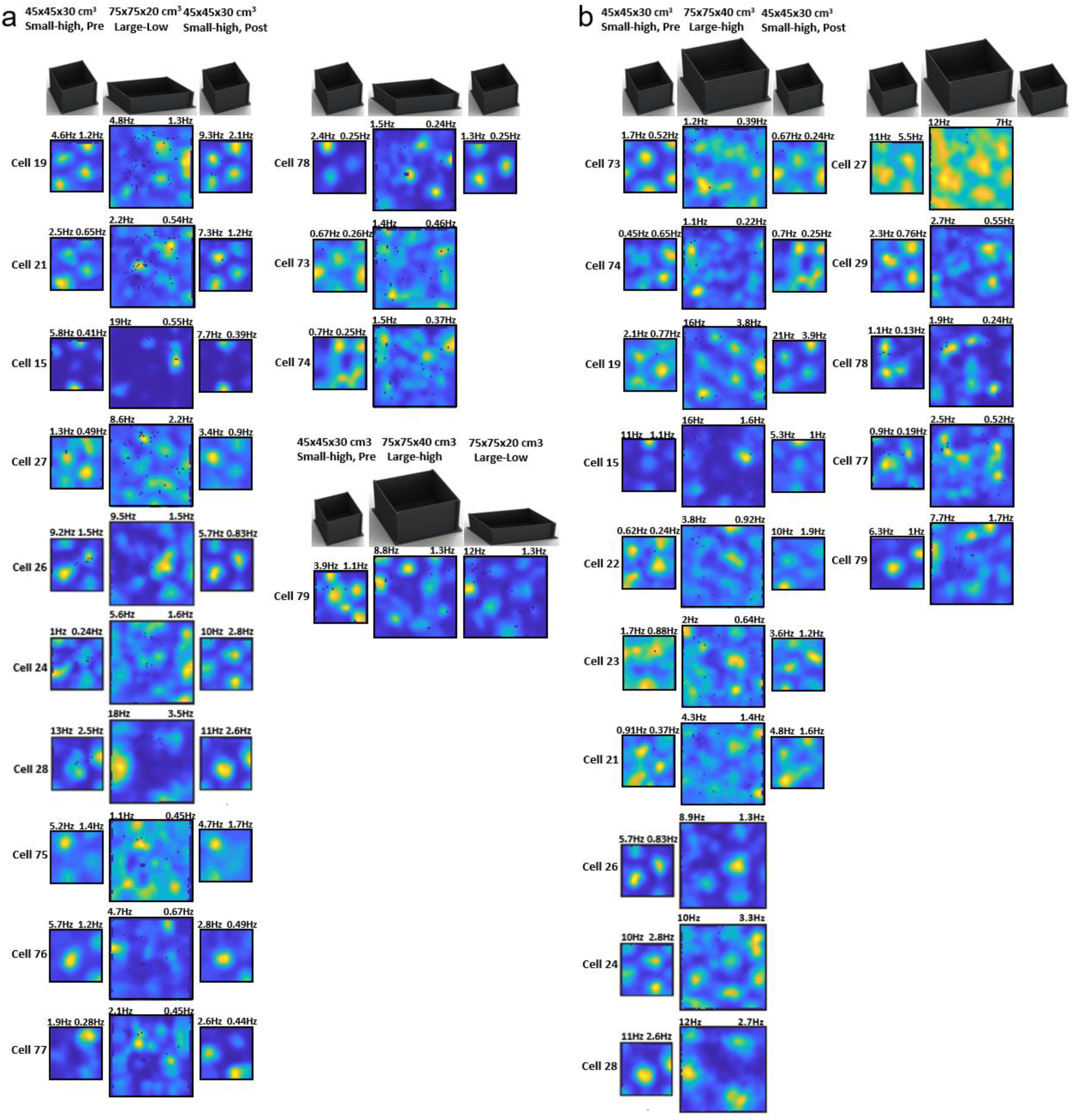
Firing rate maps of all grid cells recorded in mice after gaining experience (Experienced) with navigation in the large-scaled arena. Mice explored different sets of sequences of arenas within one recording day. Arenas included a small-size arena with high walls (45×45×30 cm^3^, Small-high, pre) and a large-size arena with either **a)** low walls (75×75×20 cm^3^, Large-low) or **b)** high walls (75×75×40 cm^3^, Large-high). The numbers above the map show peak and mean rates. Yellow and blue colors indicate high and low values, respectively.

**Extended Data Fig. 7:**
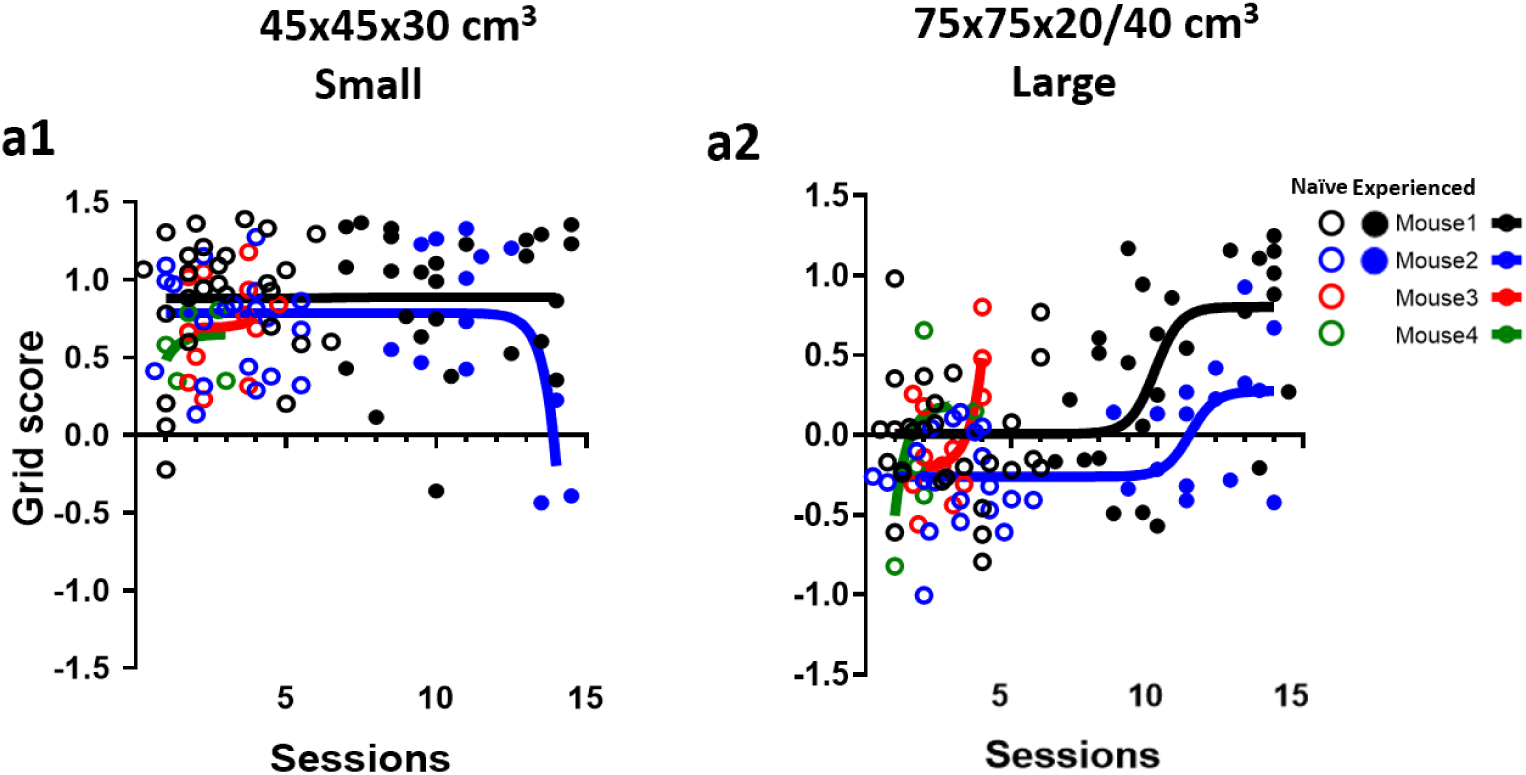
Data on changes in grid scores with increasing number of recording sessions performed on separate days. **a1** shows data on changes in grid scores when mice explored a small-size arena with high walls. **a2** shows data on changes in grid scores associated when mice explored a large-scaled arena with either low or high walls. **a1 and a2**, Open circles show data on naïve mice and filled circles show data on experienced mice. Different circle colors indicate data from a total of four different mice.

**Extended Data Table 1.** Spatial periodicity in grid cell firing is disrupted in largely scaled environments.

|  |  |  |  |  |  |  |  |  |
| --- | --- | --- | --- | --- | --- | --- | --- | --- |
| **Fig. 1c** | Mixed model fixed effect (type III) |  | P value |  | F (DFn, DFd) |  |  |  |
|  | Treatment (between columns) |  | <0.0001 |  | F (1.476, 35.42) = 71.14 |  |  |  |
|  | Tukey's multiple comparisons test | Mean Diff. | 95.00% CI of diff. | P Value | SE of diff. | n1 | n2 | DF |
|  | 45x45x30 Pre vs. 75x75x20 | 1.056 | 0.7950 to 1.317 | <0.0001 | 0.1035 | 22 | 22 | 21 |
|  | 45x45x30 Pre vs. 45x45x30 Post | 0.1203 | -0.1159 to 0.3566 | 0.4156 | 0.09299 | 20 | 20 | 19 |
|  | 75x75x20 vs. 45x45x30 Post | -0.9247 | -1.215 to -0.6342 | <0.0001 | 0.1172 | 28 | 28 | 27 |
| **Fig. 1d** | Mixed model fixed effect (type III) |  | P value |  | F (DFn, DFd) |  |  |  |
|  | Treatment (between columns) |  | <0.0001 |  | F (1.586, 26.16) = 45.11 |  |  |  |
|  | Tukey's multiple comparisons test | Mean Diff. | 95.00% CI of diff. | P Value | SE of diff. | n1 | n2 | DF |
|  | 45x45x30 Pre vs. 75x75x40 | 0.691 | 0.3467 to 1.035 | 0.0003 | 0.1326 | 16 | 16 | 15 |
|  | 45x45x30 Pre vs. 45x45x30 Post | -0.1728 | -0.3986 to 0.05311 | 0.1497 | 0.08696 | 16 | 16 | 15 |
|  | 75x75x40 vs. 45x45x30 Post | -0.8577 | -1.075 to -0.6407 | <0.0001 | 0.08503 | 19 | 19 | 18 |
| **Fig. 1e** | Mixed model fixed effect (type III) |  | P value |  | F (DFn, DFd) |  |  |  |
|  | Treatment (between columns) |  | 0.0118 |  | F (1.116, 15.63) = 7.717 |  |  |  |
|  | Tukey's multiple comparisons test | Mean Diff. | 95.00% CI of diff. | P Value | SE of diff. | n1 | n2 | DF |
|  | 45x45x30 Pre vs. 45x45x12 | 0.4761 | 0.1663 to 0.8184 | 0.0127 | 0.1469 | 18 | 18 | 17 |
|  | 45x45x30 Pre vs. 45x45x30 Post | 0.06275 | -0.3072 to 0.1235 | 0.5652 | 0.06002 | 12 | 12 | 11 |
|  | 45x45x12 vs. 45x45x30 Post | -0.2426 | -1.098 to -0.07013 | 0.3998 | 0.18 | 12 | 12 | 11 |
| **Fig. 1g** | Wilcoxon matched-pairs Signed Rank Test | Sum of signed ranks (W) | Sum of positive, negative ranks |  | P value | n1 | n2 |  |
|  | 45x45x30 vs. 75x75x20 | -208 | 1.000, -209.0 |  | <0.0001 | 20 | 20 |  |
|  | Wilcoxon Signed Rank Test | Sum of signed ranks (W) |  |  | P value | n |  |  |
|  | 45x45x30 | 50 |  |  | 0.3683 | 20 |  |  |
|  | 75x75x20 | -368 |  |  | <0.0001 | 28 |  |  |
| **Fig. 1h** | Wilcoxon matched-pairs Signed Rank Test | Sum of signed ranks (W) | Sum of positive, negative ranks |  | P value | n1 | n2 |  |
|  | 45x45x30 vs. 75x75x40 | -124 | 6.000, -130.0 |  | 0.0004 | 16 | 16 |  |
|  | Wilcoxon Signed Rank Test | Sum of signed ranks (W) |  |  | P value | n |  |  |
|  | 45x45x30 | -76 |  |  | 0.0507 | 16 |  |  |
|  | 75x75x40 | -190 |  |  | <0.0001 | 19 |  |  |
| **Fig. 1i** | Wilcoxon matched-pairs Signed Rank Test | Sum of signed ranks (W) | Sum of positive, negative ranks |  | P value | n1 | n2 |  |
|  | 45x45x30 vs. 45x45x12 | -23 | 27.00, -51.00 |  | 0.3804 | 12 | 12 |  |
|  | Wilcoxon Signed Rank Test | Sum of signed ranks (W) |  |  | P value | n |  |  |
|  | 45x45x30 | 14 |  |  | 0.6221 | 12 |  |  |
|  | 45x45x12 | -32 |  |  | 0.2334 | 12 |  |  |
| **Fig. 1j** | Kruskal-Wallis test | 11 | P value | 0.0041 |  |  |  |  |
|  | Dunn's multiple comparisons test | Mean rank diff. | P value | n1 | n2 | Z |  |  |
|  | 75x75x20 vs. 75x75x40 | -1.921 | >0.9999 | 28 | 19 | 0.3763 |  |  |
|  | 75x75x20 vs. 45x45x12 | -19.08 | 0.0038 | 28 | 12 | 3.22 |  |  |
|  | 75x75x40 vs. 45x45x12 | -17.16 | 0.0202 | 19 | 12 | 2.71 |  |  |

**Extended Data Table 2.** Spatial periodicity in grid cell firing is maintained in medium-scaled environments.

|  |  |  |  |  |  |  |  |  |
| --- | --- | --- | --- | --- | --- | --- | --- | --- |
| Fig. 2c | Mixed model fixed effect (type III) |  | P value |  | F (DFn, DFd) |  |  |  |
|  | Treatment (between columns) |  | 0.029 |  | F (1.507, 27.13) = 4.512 |  |  |  |
|  | Tukey's multiple comparisons test | Mean Diff. | 95.00% CI of diff. | P Value | SE of diff. | n1 | n2 | DF |
|  | 45x45x30 Pre vs. 60x60x15 | 0.2872 | -0.3105 to 0.8849 | 0.4311 | 0.224 | 13 | 13 | 12 |
|  | 45x45x30 Pre vs. 45x45x30 Post | -0.1887 | -0.6963 to 0.3188 | 0.5821 | 0.1852 | 11 | 11 | 10 |
|  | 60x60x15 vs. 45x45x30 Post | -0.4412 | -0.8254 to -0.05691 | 0.0252 | 0.1423 | 12 | 12 | 11 |
| Fig. 2d | Mixed model fixed effect (type III) |  | P value |  | F (DFn, DFd) |  |  |  |
|  | Treatment (between columns) |  | 0.083 |  | F (1.721, 24.09) = 2.870 |  |  |  |
| Fig. 2e | Kruskal-Wallis test |  | 7.261 | P value |  | 0.064 |  |  |
| Fig. 2f | Mixed model fixed effect (type III) |  | P value |  | F (DFn, DFd) |  |  |  |
|  | Treatment (between columns) |  | 0.0006 |  | F (1.705, 16.20) = 13.07 |  |  |  |
|  | Tukey's multiple comparisons test | Mean Diff. | 95.00% CI of diff. | P Value | SE of diff. | n1 | n2 | DF |
|  | 45x45x30 Pre vs. 60x60x15 | -4.293 | -7.358 to -1.228 | 0.0083 | 1.118 | 11 | 11 | 10 |
|  | 45x45x30 Pre vs. 45x45x30 Post | -0.6167 | -3.786 to 2.553 | 0.8464 | 1.109 | 9 | 9 | 8 |
|  | 60x60x15 vs. 45x45x30 Post | 5.331 | 2.155 to 8.507 | 0.0029 | 1.137 | 10 | 10 | 9 |
| Fig. 2g | Mixed model fixed effect (type III) |  | P value |  | F (DFn, DFd) |  |  |  |
|  | Treatment (between columns) |  | <0.0001 |  | F (1.272, 17.81) = 33.89 |  |  |  |
|  | Tukey's multiple comparisons test | Mean Diff. | 95.00% CI of diff. | P Value | SE of diff. | n1 | n2 | DF |
|  | 45x45x30 Pre vs. 60x60x45 | -4.466 | -6.673 to -2.259 | 0.0003 | 0.8433 | 15 | 15 | 14 |
|  | 45x45x30 Pre vs. 45x45x30 Post | 1.054 | 0.1330 to 1.974 | 0.0245 | 0.3518 | 15 | 15 | 14 |
|  | 60x60x45 vs. 45x45x30 Post | 5.519 | 3.353 to 7.686 | <0.0001 | 0.8278 | 15 | 15 | 14 |
| Fig. 2h | Kruskal-Wallis test |  | 19.52 | P value |  | 0.0002 |  |  |
|  | Dunn's multiple comparisons (two showed) |  | Mean rank diff. |  | P value | n1 | n2 | Z |
|  | 45x45x30' pre vs. 45x45x30 pre |  | -2.133 |  | >0.9999 | 13 | 15 | 0.3392 |
|  | 60x60x15' vs. 60x60x45 |  | -0.03333 |  | >0.9999 | 14 | 15 | 0.0054 |
| Fig. 2i | Mann Whitney test | Mann-Whitney U | Sum of ranks in column A,B |  | P value | n1 | n2 |  |
|  | 60x60x15 vs. 75x75x20 | 43 | 359, 421 |  | 0.0001 | 12 | 27 |  |
|  | Wilcoxon Signed Rank Test | Sum of signed ranks (W) |  |  | P value | n |  |  |
|  | 60x60x15 | 64 |  |  | 0.0093 | 12 |  |  |
|  | 75x75x20 | -268 |  |  | 0.0007 | 27 |  |  |
| Fig. 2j | Mann Whitney test | Mann-Whitney U | Sum of ranks in column A,B |  | P value | n1 | n2 |  |
|  | 60x60x45 vs. 75x75x40 | 47 | 358, 237 |  | 0.0006 | 15 | 19 |  |
|  | Wilcoxon Signed Rank Test | Sum of signed ranks (W) |  |  | P value | n |  |  |
|  | 60x60x45 | 104 |  |  | 0.0015 | 15 |  |  |
|  | 75x75x40 | -50 |  |  | 0.3321 | 19 |  |  |
| Fig. 2k | Mann Whitney test | Mann-Whitney U | Sum of ranks in column A,B |  | P value | n1 | n2 |  |
|  | 60x60x15 vs. 75x75x20 | 30 | 215, 220 |  | 0.002 | 10 | 19 |  |
|  | Wilcoxon Signed Rank Test | Sum of signed ranks (W) |  |  | P value | n |  |  |
|  | 60x60x15 | -33 |  |  | 0.1055 | 10 |  |  |
|  | 75x75x20 | -188 |  |  | <0.0001 | 19 |  |  |
| Fig. 2l | Mann Whitney test | Mann-Whitney U | Sum of ranks in column A,B |  | P value | n1 | n2 |  |
|  | 60x60x45 vs. 75x75x40 | 51 | 354, 241 |  | 0.001 | 15 | 19 |  |
|  | Wilcoxon Signed Rank Test | Sum of signed ranks (W) |  |  | P value | n |  |  |
|  | 60x60x45 | -64 |  |  | 0.073 | 15 |  |  |
|  | 75x75x40 | -190 |  |  | <0.0001 | 19 |  |  |
| Fig. 2n | Mann Whitney test | Mann-Whitney U | Sum of ranks in column A,B |  | P value | n1 | n2 |  |
|  | 45x45x30 Pre x 45x45x45 Post vs. 45x45x30 Pre x 60x60x15 | 13.5 | 79.50 , 220.5 |  | 0.0003 | 11 | 13 |  |
| Fig. 2o | Mann Whitney test | Mann-Whitney U | Sum of ranks in column A,B |  | P value | n1 | n2 |  |
|  | 45x45x30 Pre x 45x45x45 Post vs. 45x45x30 Pre x 60x60x45 | 104 | 211 , 224 |  | 0.9742 | 10 | 19 |  |

**Extended Data Table 3.** Grid maps in large-scaled environments are experience-dependent.

|  |  |  |  |  |  |  |  |  |
| --- | --- | --- | --- | --- | --- | --- | --- | --- |
| Fig. 3c | Mixed model fixed effect (type III) |  | P value |  | F (DFn, DFd) |  |  |  |
|  | Treatment (between columns) |  | 0.0174 |  | F (1.666, 19.16) = 5.432 |  |  |  |
|  | Tukey's multiple comparisons test | Mean Diff. | 95.00% CI of diff. | P Value | SE of diff. | n1 | n2 | DF |
|  | 45x45x30 Pre vs. 75x75x20 | 0.5013 | -0.06707 to 1.070 | 0.0869 | 0.2153 | 14 | 14 | 13 |
|  | 45x45x30 Pre vs. 45x45x30 Post | -0.0109 | -0.5316 to 0.5098 | 0.9982 | 0.1899 | 11 | 11 | 10 |
|  | 75x75x20 vs. 45x45x30 Post | -0.6037 | -1.058 to -0.1490 | 0.0115 | 0.1659 | 11 | 11 | 10 |
| Fig. 3d | Mixed model fixed effect (type III) |  | P value |  | F (DFn, DFd) |  |  |  |
|  | Treatment (between columns) |  | 0.0243 |  | F (1.848, 19.40) = 4.666 |  |  |  |
|  | Tukey's multiple comparisons test | Mean Diff. | 95.00% CI of diff. | P Value | SE of diff. | n1 | n2 | DF |
|  | 45x45x30 Pre vs. 75x75x40 | 0.3103 | -0.1043 to 0.7248 | 0.1592 | 0.1584 | 15 | 15 | 14 |
|  | 45x45x30 Pre vs. 45x45x30 Post | -0.1603 | -0.6096 to 0.2891 | 0.5715 | 0.1526 | 8 | 8 | 7 |
|  | 75x75x40 vs. 45x45x30 Post | -0.7005 | -1.326 to -0.07463 | 0.0312 | 0.2125 | 8 | 8 | 7 |
| Fig. 3e | Mann Whitney test | Mann-Whitney U | Sum of ranks in column A,B | P value |  | n1 | n2 |  |
|  | 75x75x20 vs. 75x75x40 | 87 | 192, 243 | 0.4509 |  | 14 | 15 |  |
|  | Wilcoxon Signed Rank Test | Sum of signed ranks (W) |  | P value |  | n |  |  |
|  | 75x75x20 Experienced | 51 |  | 0.1189 |  | 14 |  |  |
|  | 75x75x40 Experienced | 84 |  | 0.0151 |  | 15 |  |  |
| Fig. 3f | Mixed model fixed effect (type III) |  | P value |  | F (DFn, DFd) |  |  |  |
|  | Treatment (between columns) |  | 0.0425 |  | F (1.325, 18.55) = 4.359 |  |  |  |
|  | Tukey's multiple comparisons test | Mean Diff. | 95.00% CI of diff. | P Value | SE of diff. | n1 | n2 | DF |
|  | 45x45x30 Pre vs. 75x75x20 | -5.174 | -10.57 to 0.2224 | 0.0608 | 2.044 | 14 | 14 | 13 |
|  | 45x45x30 Pre vs. 45x45x30 Post | -0.9447 | -2.951 to 1.061 | 0.4224 | 0.7184 | 10 | 10 | 9 |
|  | 75x75x20 vs. 45x45x30 Post | 2.441 | -1.323 to 6.205 | 0.2204 | 1.348 | 10 | 10 | 9 |
| Fig. 3g | Mixed model fixed effect (type III) |  | P value |  | F (DFn, DFd) |  |  |  |
|  | Treatment (between columns) |  | 0.0117 |  | F (1.106, 11.62) = 8.547 |  |  |  |
|  | Tukey's multiple comparisons test | Mean Diff. | 95.00% CI of diff. | P Value | SE of diff. | n1 | n2 | DF |
|  | 45x45x30 Pre vs. 75x75x40 | -4.769 | -8.951 to -0.5874 | 0.025 | 1.598 | 15 | 15 | 14 |
|  | 45x45x30 Pre vs. 45x45x30 Post | 1.963 | -0.7120 to 4.638 | 0.1466 | 0.9083 | 8 | 8 | 7 |
|  | 75x75x40 vs. 45x45x30 Post | 5.148 | -0.8419 to 11.14 | 0.0882 | 2.034 | 8 | 8 | 7 |
| Fig. 3h | Mann Whitney test | Mann-Whitney U | Sum of ranks in column A,B | P value |  | n1 | n2 |  |
|  | 75x75x20 Naïve vs. 75x75x20 Experienced | 82 | 547 , 443 | 0.0009 |  | 30 | 14 |  |
| Fig. 3i | Mann Whitney test | Mann-Whitney U | Sum of ranks in column A,B | P value |  | n1 | n2 |  |
|  | 75x75x40 Naïve vs. 75x75x40 Experienced | 62 | 252, 343 | 0.0045 |  | 19 | 15 |  |
| Fig. 3j | Wilcoxon matched-pairs signed rank test |  | Sum of signed ranks (W) |  | P value |  | n1 | n2 |
|  | 60x60x15 Naïve vs. 45x45x30 Post |  | -66 |  | 0.001 |  | 13 | 11 |
|  | 60x60x45 Naïve vs. 45x45x30 Post |  | -116 |  | 0.0002 |  | 15 | 15 |
|  | 75x75x20 Experienced vs. 45x45x30 Post |  | -31 |  | 0.1309 |  | 14 | 10 |
|  | 75x75x40 Experienced vs. 45x45x30 Post |  | -28 |  | 0.0547 |  | 15 | 8 |
|  | Wilcoxon Signed Rank Test |  | Sum of signed ranks (W) |  | P value |  | n |  |
|  | 60x60x15 Naïve vs. 45x45x30 Pre |  | 91 \| 22 |  | 0.0002 \| 0.3652 |  | 13 \| 11 |  |
|  | 60x60x45 Naïve vs. 45x45x30 Pre |  | 108 \| -84 |  | 0.0009 \| 0.0151 |  | 15 \| 15 |  |
|  | 75x75x20 Experienced vs. 45x45x30 Pre |  | 73 \| 23 |  | 0.0203 \| 0.2754 |  | 14 \| 10 |  |
|  | 75x75x40 Experienced vs. 45x45x30 Pre |  | 96 \| -26 |  | 0.0043 \| 0.0787 |  | 15 \| 8 |  |
| Fig. 3k | Best-fit values |  | 45x45x30 |  | 75x75x20/40 |  |  |  |
|  | Bottom |  | 0.7722 |  | -0.1142 |  |  |  |
|  | Top |  | 0.8181 |  | 0.3501 |  |  |  |
|  | LogEC50 |  | 6.037 |  | 7.411 |  |  |  |
|  | 95% CI (profile likelihood) |  |  |  |  |  |  |  |
|  | Bottom |  | undefined to 0.8795 |  | -0.2487 to 0.06958 |  |  |  |
|  | Top |  | 0.6838 to 0.9546 |  | 0.1867 to 1.109 |  |  |  |
|  | LogEC50 |  | undefined |  | 4.974 to 13.53 |  |  |  |
|  | Goodness of Fit |  |  |  |  |  |  |  |
|  | Degrees of Freedom |  | 101 |  | 104 |  |  |  |
|  | R squared |  | 0.002644 |  | 0.1913 |  |  |  |
|  | Sum of Squares |  | 18.27 |  | 21.32 |  |  |  |
|  | Sy.x |  | 0.4253 |  | 0.4525 |  |  |  |

